# Optimizing an avian influenza vaccine using a novel Bacterial Enzymatic Combinatorial Chemistry (BECC) TLR4 adjuvant

**DOI:** 10.64898/2026.03.03.709477

**Authors:** Devon V. Riley, Lauren Baracco, Sayan Das, Brandon M. Tenaglia, Sydney Speed, Carly Dillen, Juliahna Hayes, Samanta Del Veliz, Haye Nijhuis, Valerie Le Sage, Lynda Coughlan, Weina Sun, Matthew B. Frieman, Robert K. Ernst

**Affiliations:** Department of Microbiology and Immunology, School of Medicine, University of Maryland, Baltimore, MD, USA; Department of Microbial Pathogenesis, School of Dentistry, University of Maryland, Baltimore, MD, USA; Center for Vaccine Research, University of Pittsburgh School of Medicine, Pittsburgh, PA, USA; Department of Microbiology and Molecular Genetics, University of Pittsburgh School of Medicine, Pittsburgh, PA, USA; Center for Vaccine Development and Global Health (CVD), University of Maryland School of Medicine, Baltimore, MD, USA; Department of Microbiology at Icahn School of Medicine at Mount Sinai, New York, NY, USA

## Abstract

The development of broadly protective and dose-sparing influenza vaccines remains a critical challenge, particularly for zoonotic H5N1 strains with pandemic potential. This study evaluates BECC470s, a synthetic TLR4 adjuvant, for its ability to enhance the immunogenicity and protective efficacy of recombinant H5 hemagglutinin (rHA) vaccination in murine models. BECC470s-adjuvanted rHA elicited robust IgG1/IgG2a antibody responses and complete survival following homologous 2004 H5N1 challenge in a prime–boost model. Although BECC470s broadened antibody binding to both variable HA head and conserved stalk domains by ELISA, functional neutralizing antibody responses were restricted to the matched 2004 H5N1 isolate, with no detectable neutralization of H5N1 viruses isolated in 2022 or 2024. These data indicate that BECC470s enhances the magnitude and apparent breadth of binding antibody responses while maintaining strain-specific neutralizing activity, supporting its potential as an adjuvant for next-generation influenza vaccines while underscoring the need for further optimization to achieve true cross-neutralizing protection.

## Introduction

Influenza virus infection remains a major global public health challenge. In the U.S., between 2010 and 2024, the CDC estimates that influenza has led to approximately 9.3 million to 41 million illnesses, 120,000 to 710,000 hospital admissions, and between 6,300 and 52,000 deaths annually (https://www.cdc.gov/flu-burden/php/about/index.html). The resulting hospitalizations and morbidity imposed a substantial economic burden through both lost productivity and increased healthcare costs. While annual influenza epidemics are primarily driven by seasonal influenza A and B viruses, highly pathogenic avian influenza (HPAI) viruses, such as H5 and H7, occasionally infect humans, occasionally infect humans and continue to pose a persistent zoonotic and pandemic threat. These viruses are particularly concerning due to their potential to cause severe disease, with mortality rates exceeding 50% during some outbreaks (https://www.cdc.gov/bird-flu/situation-summary/wildbirds.html). Developing effective human avian influenza virus vaccines is essential to address this gap and reduce the risks associated with emerging zoonotic and pandemic strains. Current vaccines for avian influenza virus in humans, especially those based on the H5 subtype of the hemagglutinin (HA) protein, have substantial limitations. These vaccines frequently exhibit low immunogenicity, requiring large antigen doses (up to 90 µg per dose) or multiple booster immunizations to elicit protective antibody responses^1–3^ due to their poor recognition by the human immune system^4,5^.

Although antibody responses typically focus on the HA head, generating antibodies directed toward both the HA head and stalk domains of influenza A virus (IAV), along with targeting key antigenic sites on H5 HA, is critical due to their complementary roles in virus neutralization and cross-protection. HA1 contains the globular head domain, which is highly variable between subtypes, and HA2, which consists of stalk and a small part of HA1^6^ and is more conserved between subtypes. Antibodies directed against the HA head domain inhibit the interaction between viral particles and sialic acid receptors^7^, thereby playing a pivotal role in the neutralization of specific influenza virus strains; whereas those recognizing the conserved HA stalk domain contribute to broad cross-protective immunity, underscoring the necessity to generate both antibody subsets in vaccination strategies.

Adjuvants offer a promising strategy to overcome these limitations by not only enhancing the magnitude of the immune response but also by modulating its qualitative aspects by shaping the nature and durability of the immune response through immunomodulatory mechanisms^8^. In particular, adjuvants that activate innate immune receptors, such as Toll-like receptors (TLRs) can drive potent and durable humoral and cellular immunity^9,10^. BECC470s is a rationally engineered synthetic Toll-like receptor 4 (TLR4) ligand derived through bacterial enzymatic combinatorial chemistry (BECC). It is designed to activate innate immunity by mimicking and modifying known lipid A structures, thereby eliciting a balanced and potent immune response to enhance vaccine efficacy^11^.

Building on our previous findings that BECC470s enhances immune responses to trimeric H1 HA^12–14^, our current results demonstrate that BECC470s is a potent adjuvant that can also significantly enhances the immunogenicity of the H5 HA antigen, resulting in improved antibody responses and increased protective efficacy at low vaccine doses. Here, we report that BECC470s augments the protective efficacy of a recombinantly produced H5 HA (rHA) derived from A/Vietnam/1203/2004 (H5N1) by enhancing humoral responses and broadening epitope recognition. Using a murine challenge model, mice that received a single prime immunization of BECC470-adjuvanted rHA were effectively protected against IAV-associated morbidity and mortality following a homologous IAV challenge with a 6:2 PR8 reassortant virus with low pathogenicity HA (polybasic cleavage site removed: HALo) from the A/Vietnam/1203/2004 strain^15^ (hereafter referred to as PR8/H5). Mechanistically, BECC470s broadened HA domain targeting and potentiated neutralizing antibody responses, with a notable increase in linear B cell epitope-specific antibodies focused on the mapped antigenic site B on the H5 HA protein. In addition, vaccination with H5 HA and BECC470s induced antibodies that bound a wide range of HAs from phylogenetically divergent influenza virus strains. These findings demonstrate the capacity of BECC470s to enhance both the breadth and potency of cross-reactive antibody responses essential for improved influenza vaccination strategies. Given the continuous emergence of novel influenza strains, the development of adjuvanted vaccines capable of eliciting broad and robust protection is imperative for pandemic preparedness and global health security.

## Results

### BECC470s adjuvant enhances rHA efficacy, allowing for reduced antigen doses while offering improved protection against homologous IAV challenges

We investigated the immune response and protection from challenge when mice were vaccinated with rHA in combination with BECC470s or PHAD™ (Phosphorylated Hexaacyl Disaccharide). PHAD is structurally similar to detoxified monophosphoryl lipid A (MPLA)^16^. In this study, PHAD is used as a comparator adjuvant because it is a synthetic TLR4 agonist that is structurally similar to MPLA and recapitulates its immunostimulatory properties. By directly comparing BECC470 to PHAD, we benchmark our novel TLR4-based adjuvant against a well-characterized MPLA mimetic, enabling meaningful interpretation of relative potency and immune polarization. This choice is clinically relevant, as MPLA-like TLR4 agonists are components of licensed human vaccine adjuvant systems^17,18^, so demonstrating comparable or improved performance of BECC470 versus PHAD supports the translational potential of BECC470 as a candidate adjuvant for future vaccines. To determine the minimum concentration of rHA required to confer protection against homologous IAV challenge, a concentration-response experiment was conducted using a prime-boost vaccination regimen (Day 0 and 14) with or without adjuvants. Sera was collected at day 28 post vaccination, and immunogenicity was assessed by quantifying total IgG, IgG1, and IgG2a titers via ELISA (Figs. 1A–C). BECC470s demonstrated antigen-sparing, as total IgG responses were similar between 0.05 µg and 0.1 µg rHA doses; however, the 0.1 µg rHA + BECC470s group elicited significantly higher IgG levels than 0.1 µg rHA administered without adjuvant (*p<0.0117; Fig. 1A). In mice receiving BECC470s as an adjuvant, IgG1 titers reached saturation at both concentrations, whereas PHAD required higher rHA doses to achieve comparable levels. BECC470s-adjuvanted mice exhibited significantly higher IgG1 titers than those receiving unadjuvanted vaccine or low-dose rHA with PHAD (**p < 0.08; Fig. 1B). Notably, only the 0.1 µg rHA + BECC470s group elicited significantly elevated IgG2a titers relative to all other groups (*p < 0.02), indicative of a more balanced Th1/Th2 response (Fig. 1C).

**Fig. 1.**
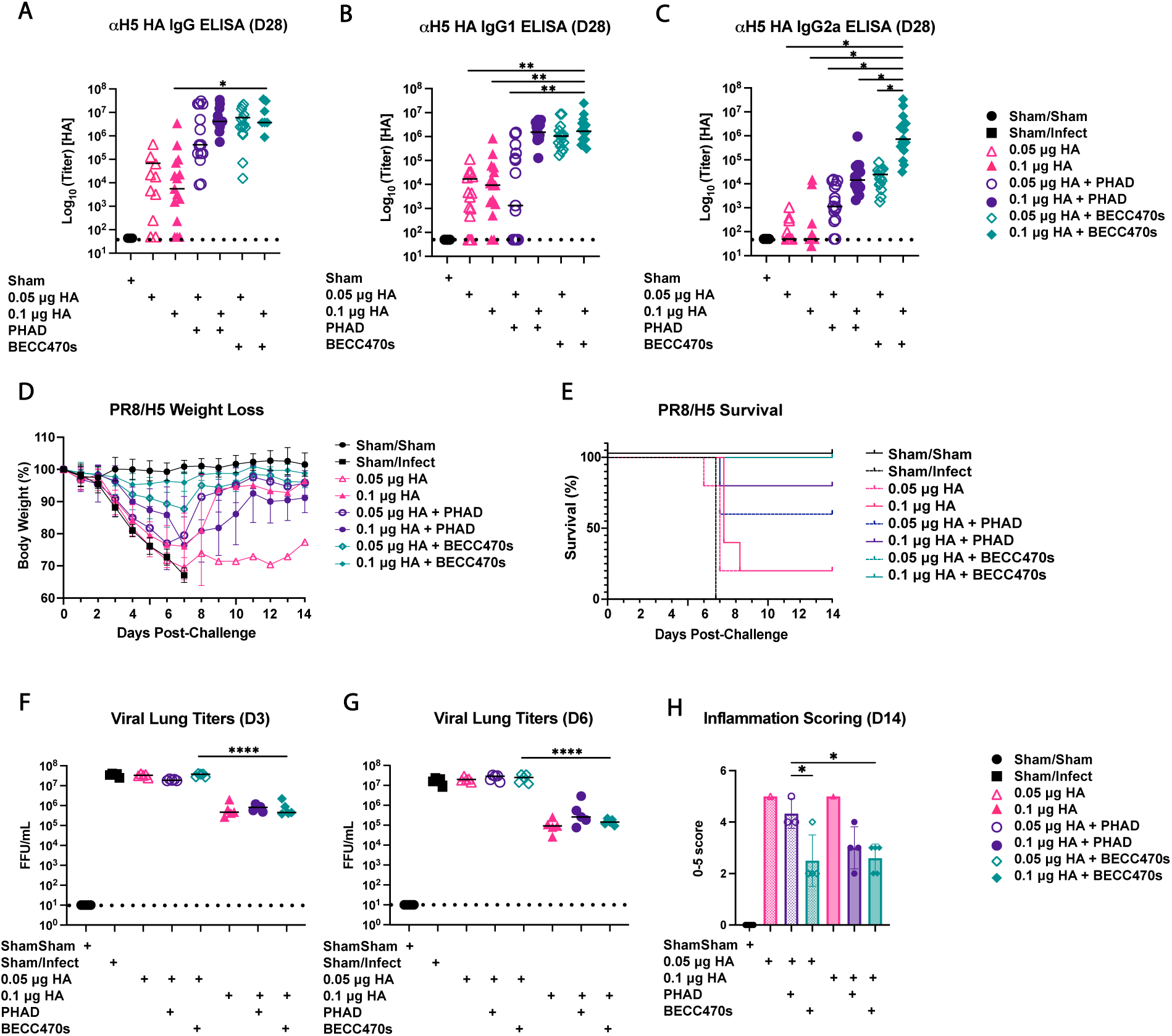
Determining rHA dose needed in a prime-boost vaccination model to protection against homologous IAV challenge. Six-week-old BALB/c mice (n=15) were immunized via a prime-boost regimen (Day 0 and 14) with 0.1 μg rHA derived from A/Vietnam/1203/2004, formulated either without adjuvant, with 50 μg PHAD, or with 50 μg BECC470s preceding PR8/H5 challenge (500 PFU) at Day 28. **A**) Pre-infection day 28 serum ELISA total IgG (*p<0.0117), IgG1 (**B**) and IgG2a (**C**) (*p<0.02, **p < 0.008). **D**)14 day weight loss in BALB/c mice (n=15) and **E**) 14 day survival graph. **F**) Virus titer of lung homogenate 3 days post infection and **G**) 6 days post infection. **H**) Averaged alveolar and peribronchiolar scoring of lung histology slides 14 days post-infection. The horizontal dashed line denotes the assay limit of detection (LOD). If not indicated otherwise, differences are not statistically significant.

On day 28, vaccinated and sham control BALB/c mice (n=15 per group) were challenged intranasally with 500 PFU of PR8/H5 and monitored daily for weight loss (Fig. 1D). Survival was assessed over 14 days post-challenge (Fig. 1E). Both unadjuvanted rHA doses failed to protect mice from weight loss, with only 20% survival in these groups. In comparison, formulations containing BECC470s provided nearly complete protection against weight loss, and the group receiving 0.1 µg rHA with BECC470s achieved total survival (Fig. 1E). Viral titers in lung homogenates collected on days 3 and 6 post-challenge were significantly reduced in the group vaccinated with 0.1 µg rHA compared to the 0.05 µg rHA group (****p<0.0001) (Fig. 1F and G). Histological analysis of lung tissues stained with Hematoxylin and Eosin (H&E) to visualize tissue structure at day 14 post-challenge revealed significant differences in both peribronchiolar and alveolar inflammation among treatment groups. While the 0.05 µg and 0.1 µg rHA vaccine groups exhibited bronchiolar and periarterial inflammation, both BECC470s-adjuvanted groups showed significantly lower inflammation scores (*p < 0.05) compared to the 0.05 µg rHA + PHAD group. Despite similar lung viral titers observed between the 0.05 µg and 0.1 µg rHA groups, inflammation at day 14 differed markedly, with the BECC470s-adjuvanted groups demonstrating the lowest inflammatory cell infiltration, with scores approaching baseline levels (Fig. 1H). Mice receiving PHAD as an adjuvant exhibited intermediate inflammation levels. Collectively, vaccination with 0.1 µg rHA combined with BECC470 significantly mitigated virus-induced lung inflammation, indicating enhanced protective potential relative to alternative vaccine formulations. These results demonstrate that BECC470s is a highly effective adjuvant that enhances immune responses, supports antigen dose sparing, and provides complete protection against homologous IAV challenge in a prime-boost vaccination model.

### BECC470s-adjuvanted rHA vaccine protects mice following a single prime dose

In the United States, FDA-approved annual influenza vaccines are administered as a single dose for adults or 1 to 2 doses for pediatric patients and are formulated as trivalent preparations containing three distinct hemagglutinin (HA) antigens (https://www.fda.gov/vaccines-blood-biologics/influenza-vaccine-composition-2025-2026-us-influenza-season). The standard adult dose includes 15 µg of HA per viral strain in the vaccine (https://www.cdc.gov/flu/professionals/acip/app/dosage.htm). Ideally, a single immunization would elicit robust titers of influenza-specific antibodies that confer protective immunity. A single-dose immunization capable of inducing strong immunity is highly advantageous during a pandemic, as it enables rapid population coverage, enhances vaccine uptake, and reduces financial and logistical burdens. With repeated vaccination, immune boosting effects occur with each dose as memory B cells recall previous exposures; however, after initial vaccination, older adults generally produce only modest levels of antibodies^19–21^.

To evaluate the efficacy of a prime-only adjuvanted vaccine against homologous influenza A virus (IAV) challenge, BALB/c mice (n=15 per group) were immunized with 0.1 or 1 µg of recombinant HA (rHA), alone or in combination with 50 µg of PHAD or BECC470s. A dose of 0.1 μg rHA combined with BECC470s was previously established as protective in a prime-boost model (Fig. 1). Antibody responses were assessed in each vaccination group to correlate with observed *in vivo* protection. Total serum IgG levels were quantified by ELISA at day 28 following immunization. The mean antibody titer for mice given 0.1 µg rHA protein alone was at the limit of detection, while those receiving 1 µg rHA alone exhibited a mean titer of approximately 10^4^. Total serum IgG measured showed significantly elevated titers in the 1 µg + BECC470s group compared to all other groups, including PHAD-containing formulation (*p<0.05, **p<0.005; Fig. 2A).

**Fig. 2.**
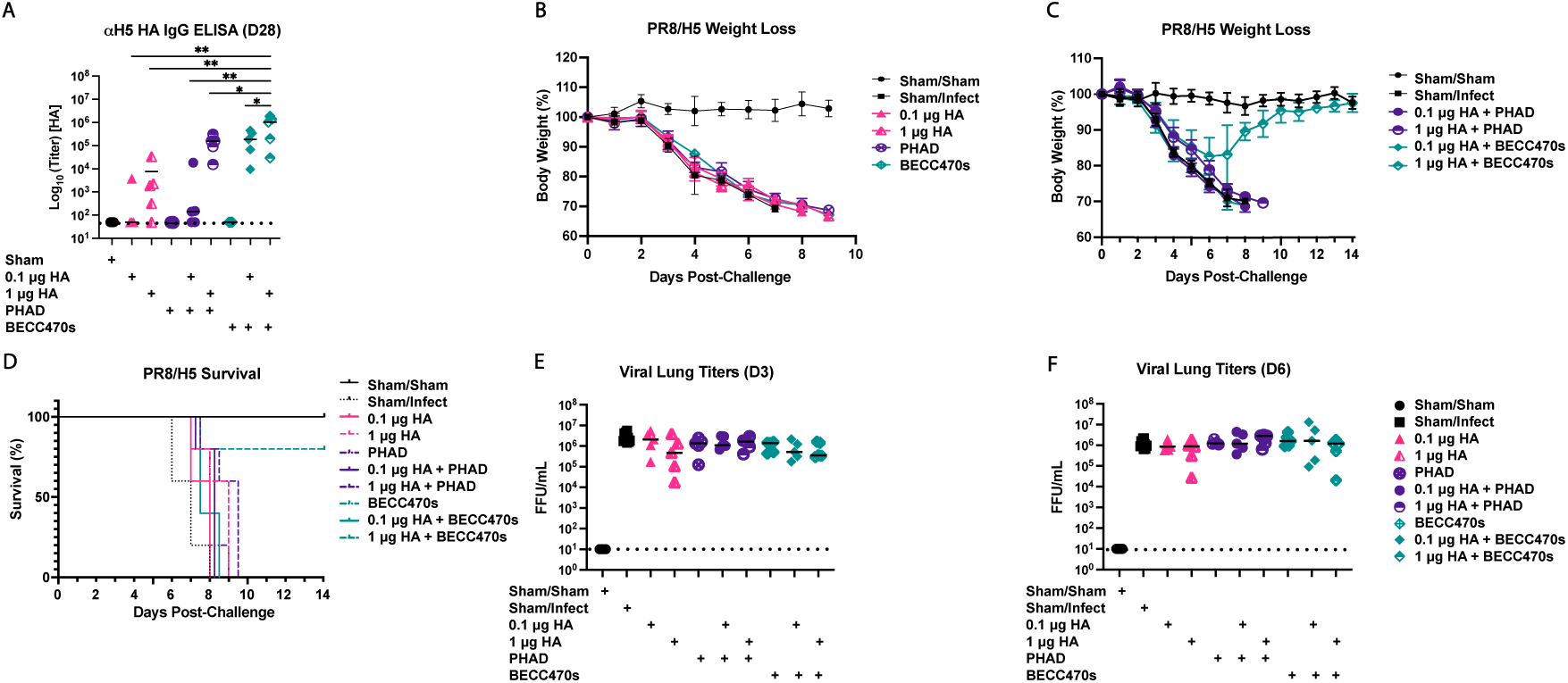
Prime-only BECC470s-adjuvanted rHA vaccine confers protection in mice. **A**) Pre-infection day 28 serum ELISA of total IgG with vaccination including 0.1 or 1 μg HA protein in combination with 50 μg of BECC470s or PHAD adjuvants in prime only vaccination schedule (*p<0.05, **p < 0.005). **B**) and **C**) 14 day weight loss in BALB/c mice (15 per group) after infection with 500 PFU of PR8/H5 with **D**) 14 day survival graph. **E**) Virus titer of lung homogenate 3 days post infection and **F**) 6 days post infection. The horizontal dashed line denotes the assay limit of detection (LOD). If not indicated otherwise, differences are not statistically significant.

Mice receiving unadjuvanted rHA or adjuvant alone showed no protection from weight loss post-challenge, losing ∼30% of their starting weight (Fig. 2B). Only the 1 µg rHA + BECC470s group demonstrated protection from ∼18% weight loss (Fig. 2C), as well as protection from mortality, with 80% survival observed in this group (Fig. 2D); all other groups required euthanasia by day 9 due to IACUC protocol regulations for weight loss.

Lung homogenates from mice collected on days 3 and 6 post-challenge were analyzed for viral load using focus forming unit (FFU) assays. Interestingly, no differences in lung viral load were observed among groups at either day 3 or day 6 post-infection (Fig. 2E, F) despite differences in clinical protection. These results demonstrate that BECC470s effectively enhances protective immunity in a prime-only influenza vaccination model, conferring significant survival benefit despite no measurable differences in viral lung load.

### BECC470s adjuvant broadens HA domain targeting and enhances the breadth and potency of cross-reactive neutralizing antibodies against influenza A

To evaluate the contribution of specific hemagglutinin regions to antibody cross-reactivity, ELISA assays were performed using antigens derived from either the HA1 domain or the conserved HA stalk. This approach allowed for a comparative analysis of antibody responses specific to each domain, thereby clarifying their respective roles in cross-reactive recognition. To determine if adjuvanting H5 HA with BECC470s increased cross-protective immunity, we analyzed the HA1 and HA stalk binding capability of antibodies from rHA vaccinated mice vaccinated with either protein alone, rHA + PHAD, or rHA+ BECC470s and showed that serum antibodies from mice receiving a prime-boost regimen demonstrated that addition of BECC470s as an adjuvant enhanced HA-specific humoral responses. Specifically, BECC470s-adjuvanted vaccines elicited significantly greater titers of both αH5 HA1 (Fig. 3A) and αH5 stalk (Fig. 3E) total IgG by day 28 post-prime compared to non-adjuvanted and PHAD groups (*p < 0.05). These elevated titers were predominantly due to increased IgG1 subclass responses (Figs. 3B and 3F), while IgG2a responses were generally lower and did not differ significantly among groups (Figs. 3C and 3G). Vaccination with rHA alone or with PHAD resulted in minimal stalk-binding antibodies, whereas BECC470s conferred a robust IgG1 response specific to the conserved HA stalk domain.

**Fig. 3.**
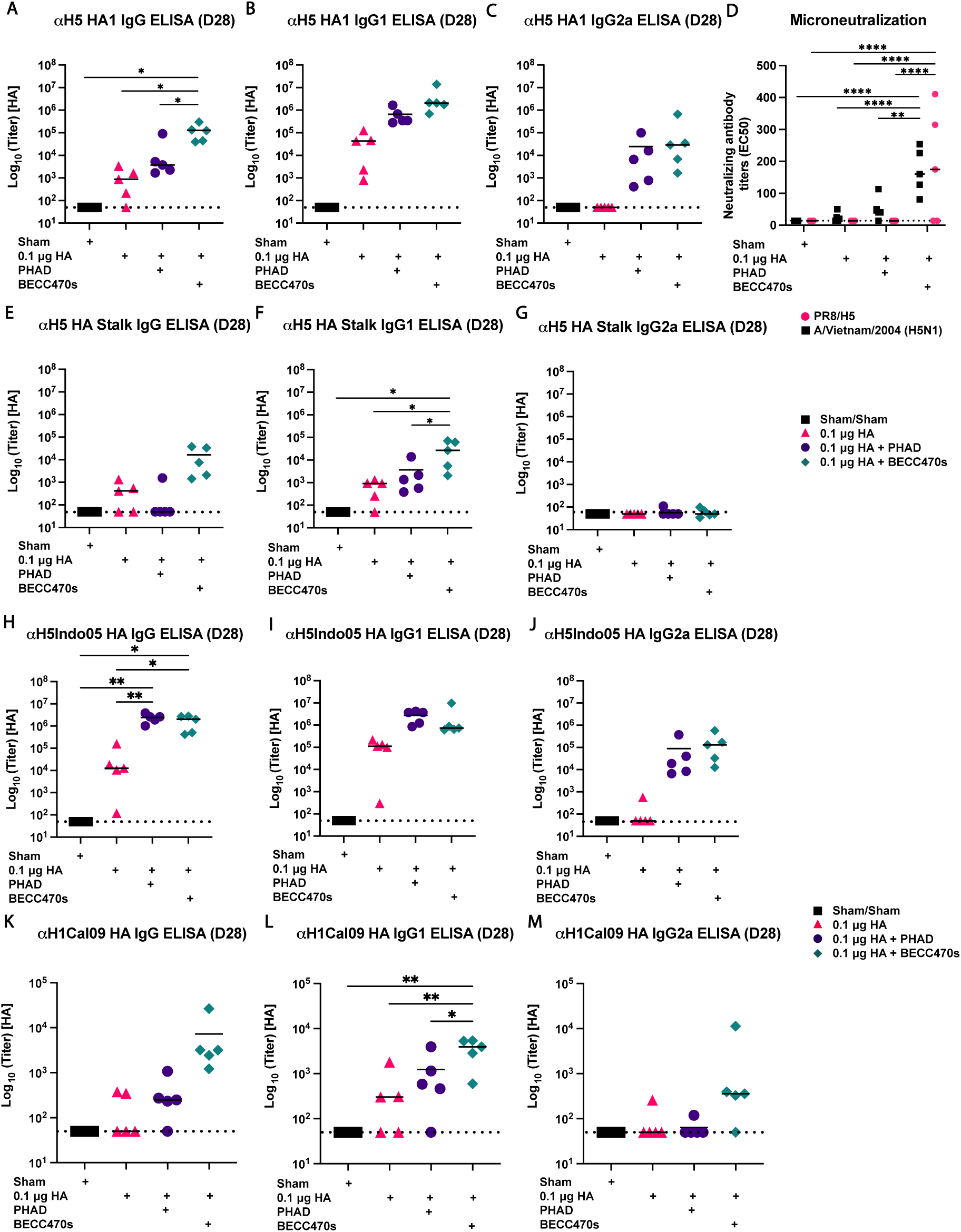
HA1, stalk-specific, neutralizing, and cross-reactive antibody responses elicited by H5 rHA vaccination in adult mice. Six-week-old BALB/c mice were immunized via a prime-boost regimen with 0.1 μg rHA derived from A/Vietnam/1203/2004, formulated either without adjuvant, with 50 μg PHAD, or with 50 μg BECC470s. Day 28 post-immunization sera was collected and antibody titers specific for αH5 HA1 were quantified by ELISA; total IgG (**A**), IgG1 (**B**), and IgG2a (**C**) levels were measured(*p < 0.05). (**D**) Neutralizing activity was evaluated by microneutralization assay against PR8/H5 (pink circles) and A/Vietnam/2004 (H5N1; black squares) viruses. The horizontal dashed line denotes the assay limit of detection (LOD). Antibody titers specific for αH5 stalk were quantified by ELISA; total IgG (**E**), IgG1 (**F**), and IgG2a (**G**) levels were measured (*p < 0.05). Antibody titers specific for αH5 or H1 HA were quantified by ELISA using A/Indonesia/5/2005 HA (**H-J**) or A/California/07/2009 HA (**K-M**) as the coating antigen. Total IgG, IgG1 and IgG2a were measured for each coating antigen. (*p < 0.02, **p < 0.009, ****p<0.0001). Prism 10 was used for statistical comparisons between using two-way ANOVA with multiple comparisons. If not indicated otherwise, differences are not statistically significant.

Binding antibodies, quantified by ELISA above, demonstrate immunogenicity but do not necessarily indicate protective efficacy. The gold-standard correlate for protection is the generation of neutralizing antibodies, as these directly inhibit viral entry and replication in host cells^22,23^. The ability of neutralizing antibodies to target influenza virus surface proteins, such as hemagglutinin, plays a fundamental role in preventing infection and limiting disease severity, underscoring their significance in both natural immunity and vaccine-induced protection^24,25^. Microneutralization (MN) assays were performed to examine adjuvant-dependent enhancement of neutralizing antibody responses using microneutralization assays against PR8/H5 (A/Vietnam/120/3/2004 HALo) or a wildtype A/Vietnam/2004 isolate (performed in BSL-3 containment). Mice immunized with BECC470s-adjuvanted H5 rHA exhibited significantly higher neutralizing antibody titers against PR8/H5 demonstrating strong inhibition for the homologous strain (****p<0.0001 compared to all other groups, Fig. 3D). To assess reactivity against H5N1 strains at BSL-3, live virus microneutralization assays were conducted against A/Vietnam/2004, as well as two recently emergent strains, A/dairy cattle/Texas/24008749001/2024 (H5N1) and A/mink/Spain/3691 8_22VIR10586-10/2022 (H5N1). While robust strain-specific neutralizing titers were observed against the original H5N1 strain in BECC470s-adjuvanted vaccinated group only (**p<0.02, ****p<0.0001 compared to all other groups, Fig. 3D), no neutralizing antibodies were detected against the cow- and mink-derived viruses (Supplementary Fig. 1). In contrast, sera from mice immunized with rHA alone, PHAD-adjuvanted vaccine, or sham controls showed neutralizing titers at or below the assay detection limit for all tested viruses. Although PHAD modestly enhanced responses relative to unadjuvanted rHA, BECC470s outperformed PHAD in eliciting potent neutralizing antibodies against homologous H5N1 viruses.

To evaluate potential cross-reactivity against more diverse HA subtypes, ELISAs against rHA from either A/Indonesia/5/2005 (H5N1) or A/California/7/2009 (H1N1) were used. The A/Indonesia/5/2005 HA shares 97% sequence homology with the immunizing A/Vietnam/1203/2005 HA, while the A/California/7/2009 HA is more divergent, sharing 63% sequence homology; all HA sequences were aligned and analyzed using Geneious Prime® version 2025.2.2.

Mice immunized with rHA alone generated low but detectable αH5 and αH1 IgG titers, whereas PHAD significantly boosted total and IgG1 responses and BECC470s further increased titers by 1–2 orders of magnitude across both specificities (*p < 0.02, **p < 0.009; Fig. 3H–M). Both adjuvants enhanced IgG1, but BECC470s consistently elicited the highest levels, and uniquely drove a marked increase in IgG2a, indicative of enhanced cross-reactive potential. These data show that BECC470s-adjuvanted H5 rHA induces robust heterologous and heterosubtypic antibody responses, supporting its promise as an adjuvant for broad protective immunity against diverse influenza A viruses.

Collectively, these findings demonstrate that BECC470s-adjuvanted H5 rHA elicits robust cross-reactive heterologous and heterosubtypic antibodies together with potent strain-specific neutralization, highlighting BECC470s as a highly effective adjuvant for broad protective humoral immunity targeting both conserved and variable HA domains across diverse influenza A virus strains.

### Enhanced breadth and magnitude of antibody responses to antigenic site B induced by BECC470s-adjuvanted rH5 HA vaccination

The broader protection observed with rHA formulated with BECC470s is hypothesized to result from both increased serum antibody levels and the induction of antibodies targeting diverse epitopes on the rHA protein. To define epitope-specific responses and quantify serum antibody magnitude post-vaccination, total IgG, IgG1, and IgG2a titers were measured against a panel of 93 overlapping peptides (12- or 17-mers with 11–amino acid overlap) spanning the H5 HA of A/Vietnam/1203/2004, following the method outlined by Das *et al*^14^. Six-week-old BALB/c mice were immunized on a prime-boost schedule with 0.1 μg rHA alone or combined with 50 μg PHAD or BECC470s. Serum samples collected on day 28 following immunization, prior to challenge, were assessed by ELISA for peptide-specific antibody binding (Fig. 4). Total IgG (Fig. 4A), IgG1 (Fig. 4B), and IgG2a (Fig. 4C) titers against individual peptide pools were determined by ELISA. Of the 93 tested peptide pools, positive antibody reactivity was detected in pool 4 (peptides 31–40) and pool 8 (peptides 72–82), with positivity defined as an O.D. value four times greater than the mean of sham-immunized controls. Pools 4 and 8 were further separated into their constituent 10 or 11 individual peptides, respectively, and assessed for antibody binding. Peptides 72–82 showed no reactivity in any group, with O.D. readings for total IgG, IgG1, and IgG2a remaining below the fourfold threshold relative to naïve controls (Supplementary Fig. 2).

**Fig. 4.**
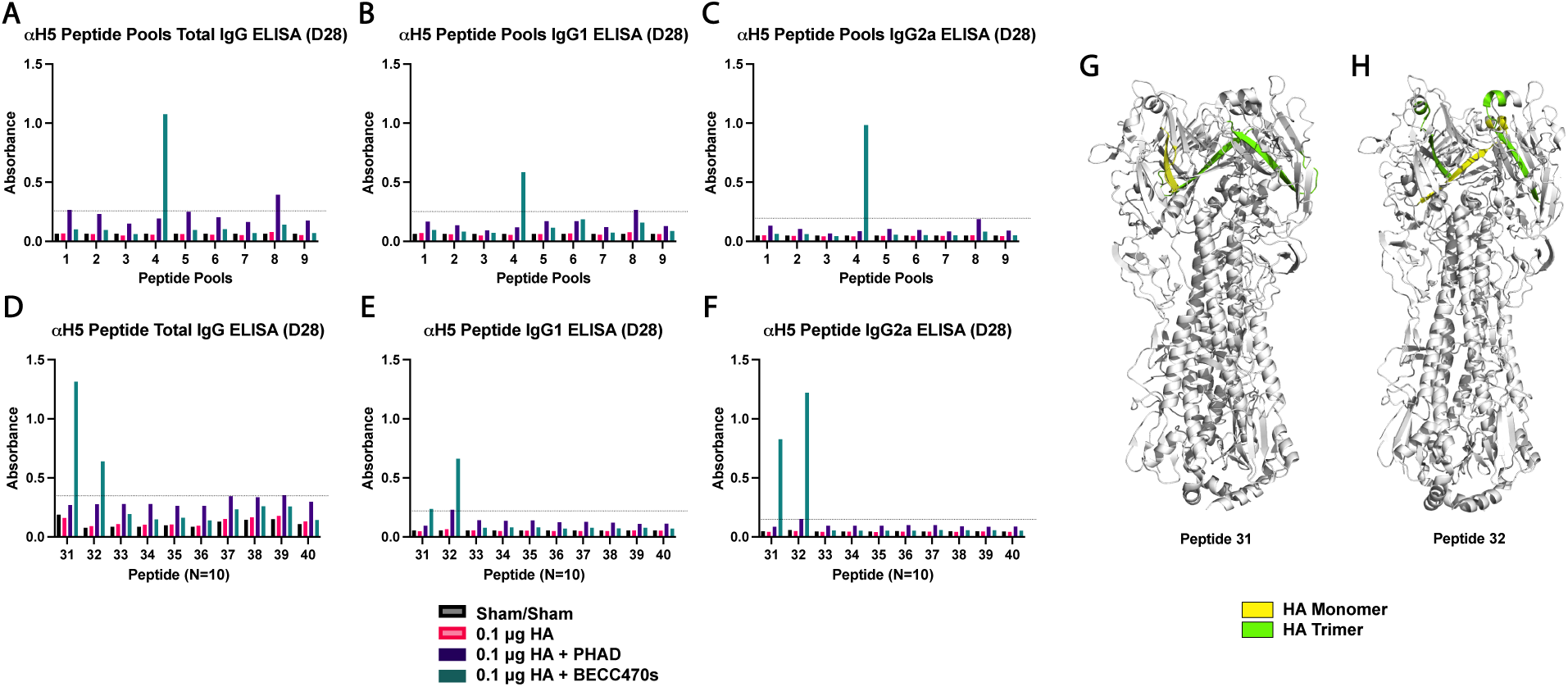
BECC470s-adjuvanted immunization broadens linear B cell epitopes. Six-week-old BALB/c mice were immunized via a prime-boost regimen with 0.1 μg rHA derived from A/Vietnam/1203/2004, formulated either without adjuvant, with 50 μg PHAD, or with 50 μg BECC470s. Initially, 93 overlapping peptides (12- or 17-mers with 11 amino acid overlaps) spanning the H5 HA of A/Vietnam/1203/2004 were pooled into nine pools. Pre-infection sera, Day 28 post-immunization, was collected and Total IgG (**A**), IgG1 (**B**) and IgG2a (**C**) titers against peptide pools were determined by ELISA. Individual peptides for pool 4 was tested with Total IgG (**D**), IgG1 (**E**) and IgG2a (**F**) determined by ELISA. The dashed line indicates an O.D. value fourfold higher than that of the sham control group; samples above this threshold are considered positive. Ribbon diagrams depicting a 3D schematic representation of peptides 31 (**G**) and 32 (**H**) created using PyMOL (HA monomer highlighted in yellow and HA trimer highlighted in green).

Mice immunized with rHA + BECC470s exhibited significantly elevated total IgG titers (Fig. 4D) against peptides 31 and 32, which correspond with antigenic site B on the HA protein^26,27^ with O.D. readings above the threshold defining positive antibody binding (dashed line). Consistent with total IgG findings, IgG subclass evaluation demonstrated that BECC470s-vaccinated mice displayed the highest IgG1 titers (Fig. 4E) toward peptides 31 and 32, reflecting a Th2-biased antibody response centered on antigenic site B. Similarly, IgG2a titers (Fig. 4F) were predominantly elevated against peptide 31, indicating a concurrent Th1-type response. In contrast, PHAD-adjuvanted and antigen-alone immunizations elicited weaker, more limited peptide-specific responses, rarely surpassing the threshold. These results demonstrate that formulation with BECC470s adjuvant significantly amplifies and broadens antibody responses targeting key linear epitopes, especially those spanning antigenic site B (peptides 31 and 32) of the H5 HA. These findings suggest that BECC470s may potentiate protective efficacy by focusing immune recognition on critical antigenic regions of the hemagglutinin protein.

## Materials and Methods

### Mouse immunogenicity and challenge study design

Mice were acclimatized for approximately five days prior to the start of experimental procedures. They were randomly assigned to cages (n = 5 per cage), with each cage representing a single experimental unit within a vaccination group. Vaccinations were administered intramuscularly (i.m.) on day 0 (prime) and day 14 (boost) using 50 μL of the assigned vaccine. Vaccine solutions were prepared by admixing 0.1 µg of rHA with a fibritin trimerization ectodomain (A/Vietnam/1203/2004) and 50 µg of either BECC470s or PHAD (Avanti, AL, USA). The vaccines were incubated with rocking for 1 hour to allow interaction between the antigen and the adjuvants. Control groups received 50 μL of sterile PBS.

Blood samples were collected at days 14 and 28 post-prime. On day 28, mice were challenged intranasally (i.n.) with PR8/H5 influenza virus at a dose of 500 PFU/mL. The PR8/H5 challenge virus was a 6:2 reassortment virus of low pathogenicity avian A/Vietnam/1203/2004, in which the polybasic cleavage site was removed (HALo), and N1 from A/Vietnam/1203/2004 in an A/Puerto Rico/8/1934 backbone. Post-challenge, mice were monitored daily for 14 days for clinical signs and weight loss. Mice exhibiting greater than 30% weight loss relative to their pre-challenge baseline were humanely euthanized by gradual CO2 exposure. All surviving mice were euthanized at day 14 post-challenge.

### Mice and Immunizations

All studies were approved by the University of Maryland School of Medicine Institutional Animal Care and Use Committee (protocol AUP-00000839) and were carried out at the University of Maryland School of Medicine Biohazard Suite. All studies were performed in accordance with the National Institutes of Health guidelines for the care and use of laboratory animals and institutional policies. All procedures complied with relevant regulations and followed the principles of the 3Rs and the ARRIVE guidelines. All mice were female BALB/cJ mice (Jackson Laboratory, Bar Harbor, ME) aged 6 to 8 weeks at the time of vaccination (n = 5–15 mice/group).

### Recombinant proteins

Recombinant HA1 and HA stalk proteins from A/Vietnam/1203/2004, as well as full-length recombinant HA (rHA) proteins from A/Vietnam/1203/2004, A/Indonesia/5/2005, A/California/7/2009, and A/BrevigMission/1/1918, were produced with C-terminal hexahistidine tag as previously described^28^. Briefly, Expi293F cells were transiently transfected with plasmids encoding the respective HA constructs and cultured in serum-free Expi expression medium following the manufacturer’s protocol. The rHA proteins used for immunization incorporated a modified transmembrane domain replaced by a heterologous carboxy-terminal trimerization motif (fibritin foldon), whereas proteins employed in ELISAs contained a GCN4 isoleucine zipper trimerization domain derived from Saccharomyces cerevisiae. Recombinant HAs were purified using NiNTA chromatography. Proper protein conformation was validated ELISA using a panel of at least 6 well-characterized monoclonal antibodies known to bind conformation-sensitive epitopes (e.g., CR9114, CT149^29,30^). Protein concentrations were quantified using NanoDrop spectrophotometry.

### ELISA

Enzyme-linked immunosorbent assays (ELISAs) were performed to quantify serum antibodies specific to recombinant hemagglutinin (rHA), following a previously described protocol with minor modifications^12,14^. Ninety-six–well plates (Nunc, NY, USA) were coated overnight at 4°C with rHA containing a GCN4 trimerization domain (2 µg/mL in 0.1 M sodium bicarbonate buffer). Plates were then washed 3 times with PBS-0.05% Tween-20 and blocked for 2 hours at room temperature with blocking buffer consisting of 0.5% nonfat dry milk and 3% goat serum in PBS-T. Serial dilutions of serum samples were added to the plates and incubated for 2 hours at room temperature. After washing, horseradish peroxidase (HRP)-conjugated secondary antibodies specific for IgG, IgG1, or IgG2a (1:3000; Thermo Scientific, MA, USA) were added to separate plates for 1 hour at room temperature. Following 3 additional washes, 100 μL of 3,3′,5,5′-Tetramethylbenzidine substrate (BD Biosciences, CA, USA) was applied, and the reaction was terminated after 5.5 minutes by adding an equal volume of stop solution (KPL, MD, USA). Antibody titers were determined using GraphPad Prism software (GraphPad Software, MA, USA).

### Peptide Microarray

Peptide array for HA (Peptide Array, Influenza Virus A/Vietnam/1203/2004 (H5N1) Hemagglutinin Protein - Influenza A virus) was obtained from BEI Resources, NIAID, NIH (Cat# NR-18974). The 93 peptide arrays (12 or 17 mers with 11 mer overlaps) were dissolved at a concentration of 1 mg/ml using manufacturer suggested solvents. 10 or 11 sequential peptides were mixed to create 9 peptide pools. Peptides were covalently linked by the free carboxylic acid group to the amino group of the 96 well plate (Thermo Scientific Covalink NH, F8 Cat#478042) by coupling agent 1-Ethyl-3-(3-dimethylaminopropyl) carbodiimide (EDAC, Sigma 341006-5GM). Briefly, pools were diluted with MES buffer (pH 6) at a final concentration of 4 ug/ml. The solution was added into 96 well plates followed by 10 μL of EDAC (10mg/ml in water). Plates were incubated overnight at room temperature and washed with distilled water. Sera from groups of mice (n=5) were pooled, and immune reactivity was assessed by ELISA as described above. To evaluate the reactivity against individual peptides in the pool, each peptide was diluted in MES buffer and coupled onto plates as described earlier.

### Determination of viral titer using focus-forming unit assay

Mouse lung tissues were homogenized in sterile PBS using a bead mill or tissue homogenizer to generate lung homogenates. Homogenates were clarified by low-speed centrifugation (1,000 x g for 10 min) and serially diluted in infection medium. Madin-Darby Canine Kidney (MDCK) cells were seeded at 5×10^4^ cells per well in 96-well plates and grown to confluence. Cells were infected with diluted lung homogenates for 1 hour at 37°C, then overlaid with infection medium containing 1% methylcellulose to restrict viral spread. After 24 hour incubation at 37°C, cells were fixed in 10% neutral buffered formalin (Sigma), quenched with 50 mM ammonium chloride, and permeabilized with 0.1% Triton X-100 in 1% BSA. After a 10-minute block with 1% BSA, cells were incubated with primary antibody (1:10,000) for 1 hour, followed by secondary antibody (1:3000) for 45 minutes. Nuclei were counterstained with Hoechst (1:3000). Wells were maintained in PBS for imaging on the Celigo cytometer. Viral foci were detected by immunostaining with a primary antibody against influenza A virus nucleoprotein, followed by a fluorophore-conjugated secondary antibody. Foci were visualized and counted using a fluorescence microscope or imaging cytometer. Virus titers were calculated as focus-forming units (FFU) per gram of lung tissue.

### Microneutralization

The microneutralization assay was performed to measure the neutralizing antibody titers against influenza A virus (IAV) using Madin-Darby Canine Kidney (MDCK) cells. Serum samples were heat-inactivated at 56°C for 30 minutes prior to use. MDCK cells were cultured in maintenance medium and trypsinized when approximately 70-90% confluent. Cells were counted and seeded at 5×10^4^ cells per well in 96-well flat-bottom plates. Sera samples in duplicate (starting dilution 1:40) were serially diluted 2-fold across the plate. Serially diluted samples were incubated with 60 µl of virus dilution (MOI of 0.078 based on FFU titers for PR8/H5) for 1 h at room temperature on a shaker. MDCK cells were then washed once with 100 µl of PBS, and 100 µl of the virus–serum mixture was added to MDCK cells. The cells were incubated in the presence of antibody and virus for 24 h at 37°C. After 24 hours cells were fixed with 10% neutral buffered formalin (Sigma). The readout was performed via immunofluorescence assay (described previously). For H5N1 virus microneutralization, cells were incubated in the presence of antibody and virus for 96 h at 37°C and readout was performed by determination of cytopathic effect. Positive virus controls (with virus only) and negative cell controls (without virus) were included on each assay plate for data normalization. Results were analyzed in Microsoft Excel and GraphPad Prism v10.6.0.

### Lung Histology

Mouse lungs were fixed by immersion in 4% paraformaldehyde (PFA) in phosphate-buffered saline (PBS) for a minimum of 48 hours. The fixed lungs were subsequently processed by the University of Maryland - Baltimore Histology Core Facility, where they were embedded in paraffin, sectioned into 5 µm slices, and stained with hematoxylin and eosin (H&E). Histological scoring was performed in a blinded manner using a scale from 0 (no inflammation) to 5 (severe inflammation), with interstitial and peribronchiolar inflammation evaluated separately. The final inflammation score was calculated by averaging these values.

### Statistical analysis

GraphPad Prism 10.6.0 was used to perform all statistical comparisons. Differences among treatment groups were analyzed as mentioned in figure legends. Exact p-values are given in figure legends, but in general a p-value of less than 0.05 was considered significant for all comparisons. * P< 0.05; ** P< 0.001.

## Discussion

Influenza A (H5N1) remains a persistent pandemic threat, necessitating the development of innovative vaccines that extend beyond the limited efficacy of current licensed formulations. Traditional influenza vaccines are often tailored to specific strains and require annual updates, leading to immunity gaps as the virus continues to evolve. The development of cross-reactive influenza vaccines is crucial for achieving broader, longer-lasting protection, minimizing the impact of viral diversity, and improving pandemic preparedness. Antigenic drift and shift in influenza viruses drive recurrent seasonal epidemics and occasional pandemics. Cross-reactive vaccines, by targeting more conserved viral components, have the potential to maintain immunity across multiple strains and epidemic seasons, reducing the impact of antigenic variability. Current FDA-approved seasonal influenza vaccines are typically trivalent, including HAs from A(H1N1), A(H3N2), and B (B/Victoria-lineage); previously, FDA-approved quadrivalent vaccines also incorporated a second B-lineage (B/Yamagata-lineage) HA^31^. While seasonal influenza vaccines are primarily designed to confer protection against circulating human strains, numerous studies have investigated their capacity to elicit cross-reactive antibodies that recognize multiple influenza A virus subtypes beyond those included in the vaccine formulation. As an example, serum from adults who received multiple annual influenza vaccinations was analyzed for cross-neutralizing activity, revealing limited neutralization of pseudovirus H5N1 strains (ranging from 36% to 0%). Antibody titers were highest for A/Japan/305/1957 (H2N2), with modest cross-neutralization against select H9 and H6 viruses. Finally, substantially less cross-reactivity toward group 2 influenza A viruses, demonstrates that current seasonal influenza vaccines do not elicit broad-spectrum neutralizing antibodies and are unlikely to confer universal protection against emerging pandemic and avian strains^32^.

Although existing FDA-approved H5N1 vaccines are safe and can stimulate immunity within their specific subtype, limitations in broad coverage and the current lack of dose-sparing strategies may present challenges for rapid, large-scale deployment in the event of a pandemic caused by a novel H5 virus^33^. Currently, only three H5N1 vaccines have received FDA approval, with two incorporating adjuvants to enhance their immunogenicity (https://www.fda.gov/vaccines-blood-biologics/vaccines/vaccines-licensed-use-united-states). Given this limited landscape, evaluating the efficacy of these available vaccines relies on established immunological endpoints. For these vaccines, the primary study endpoints to evaluate efficacy are hemagglutination inhibition (HI) antibody titers ≥ 1:40. While this is a standard goal of immunogenicity in influenza vaccine studies, HI titers do not always correlate fully with *in vivo* protection in animal models and do not measure other potential immune mechanisms like antibody-dependent cellular cytotoxicity or T cell responses^34^.

In contrast to these studies, we demonstrate that BECC470s-adjuvanted rHA vaccines markedly increase both survival and immunogenicity, as compared to control adjuvants, even with substantial antigen dose reduction or single-dose prime protocols, which is a significant advance over standard formulations. Although we did not observe statistically significant differences in lung viral titers between the adjuvanted groups at either day 3 or day 6, similar findings have been frequently reported in H5N1 studies, in which lung titers are minimally affected whereas other disease parameters including weight loss, lung pathology, or survival show more pronounced adjuvant dependent effects ^35–37^. This pattern underscores an important limitation of mouse models for evaluating H5N1 vaccines. The HA protein of IAV exhibits increased structural diversity across subtypes, driven by extensive antigenic variation within the globular head and differences in receptor-binding domains^38^. Structurally, the stalk domains of H1 and H5 HAs share conserved motifs, including the fusion peptide and F subdomain^39,40^; these regions serve as targets for broadly reactive antibodies, yet the greater divergence in HA head sequences restricts the effectiveness of cross-subtype immunity, as observed in our data. The BECC470s-adjuvanted regimen successfully amplified both the quantity and subtype breadth of antibody responses. The adjuvant-driven immune responses were marked by robust induction of both head- and stalk-directed antibodies. Notably, head-specific antibodies are potent neutralizers, yet their breadth is circumscribed. In contrast, stalk-specific antibodies engage FcγRs, inducing cellular cytotoxicity of infected cells^41^ and are essential for broader group 1 and 2 HA reactivity. Of particular interest is the enhanced targeting of antigenic site B within the HA head by BECC470s-adjuvanted formulations as measured using linear peptides. It should be noted that our binding assay selectively detects antibodies targeting linear peptide epitopes, rather than those recognizing conformational epitopes; thus, the overall magnitude and diversity of the antibody response may be underestimated.

Antigenic site B within the receptor-binding domain is a crucial target for neutralizing antibodies and antigenic drift, making it essential for immunogenicity and rational vaccine design; its immunodominance drives robust homologous protection, as evidenced by elevated neutralizing antibody titers and enhanced survival outcomes. BECC470s elicited robust neutralizing antibodies against the homologous H5N1 strain, indicating strong immunogenicity to matched antigen. The lack of neutralization against recent H5N1 isolates likely reflects antigenic divergence rather than an intrinsic limitation of BECC470s. As shown in Fig. 4, BECC470s enhances binding to antigenic site B, yet the two contemporary H5N1 strains share only ∼70% amino acid identity in this region, which may underlie the loss of neutralization. Future studies using H5 HA variants with targeted mutations in site B will be needed to test this directly. While antigenic site B is a dominant focus for immune responses, other antigenic sites on H5 HA, including sites A, C, D, and E, also play important roles in antibody recognition and viral evolution^27^. Site A, positioned within the receptor-binding domain, is a key focus for antibody binding and a hotspot for antigenic variation in H5 HA. Sites A and B display strong immunodominance, contributing substantially to antigenic drift (Supplementary Fig. 3) and facilitating viral escape from host antibody responses. Site C is situated near the head-stalk interface, overlapping regions associated with vestigial esterase and fusion functions. Additional sites, including D and E, broaden the repertoire of neutralizing antibody targets across the globular head, though they vary in exposure and structural context. This emphasizes the importance of adjuvant approaches that not only enhance immunity to these variable sites but also boost protective responses targeting conserved regions^26,27^.

We also demonstrated that BECC470s adjuvant significantly enhanced stalk-specific antibody responses against conserved HA domains, promoting broad epitope recognition across diverse influenza subtypes. Antibodies directed against the conserved stalk region of influenza hemagglutinin can protect against a wide range of viral strains, offering broad protection that could overcome the need for regular vaccine reformulation. Moreover, stalk-targeted antibodies can mediate cross-reactive immunity and contribute to the development of a universal influenza vaccine, which is believed to be key for long-lasting and broad protection against seasonal and pandemic influenza virus strains. Notably, BECC470s-adjuvanted sera demonstrated enhanced binding to both heterologous H5N1 and heterosubtypic H1N1 influenza strains, reflecting robust breadth and magnitude of the humoral immune response.

Adjuvants are well-established for enhancing the immunogenicity of influenza vaccine antigens by increasing both antibody magnitude and breadth of response, particularly in older adults and immunocompromised populations. However, the majority of seasonal influenza vaccines currently licensed for use in the United States are unadjuvanted formulations^42^. While unadjuvanted H1 influenza vaccines generally elicit stronger immune responses with 56-66% efficacy, unadjuvanted H5 vaccines tend to be poorly immunogenic, requiring higher antigen doses or multiple doses to generate measurable neutralizing antibody responses, often falling short of protective levels without adjuvant support^34,43,44^. In our prior studies examining BECC470s formulations with the H1 HA protein (admix formulation), we showed that immune responses persisted for up to 18 months after vaccination, indicating durability beyond the usual seasonal timeframe^45,46^. Additionally, BECC-adjuvanted vaccination broadened humoral targeting to include conserved HA stalk epitopes and expanded recognition of linear B-cell epitopes compared with other adjuvants, indicating improved breadth of response^14^.

Building upon BECC470s’ established immunogenicity and protective capacity in the H1 influenza model^12–14^, our findings revealed similarly robust protective outcomes with BECC470s in the H5 challenge study. Notably, adjuvanted H5 vaccine formulations achieved substantially higher efficacy than their unadjuvanted counterparts. Collectively, our results underscore BECC470s as a promising adjuvant candidate for next-generation influenza vaccines aimed at improving the breadth and durability of protection against antigenically diverse influenza viruses, addressing a major limitation of current seasonal vaccine strategies. With BECC470s’ capacity for antigen sparing and the induction of robust, well-balanced IgG1/IgG2a antibody responses, its integration into future pandemic vaccine platforms is highly advantageous.

Future vaccine platforms incorporating BECC470s may enable rapid induction of protective immunity with low antigen doses and could be optimized to broaden coverage against emerging influenza A subtypes, overcoming critical weaknesses of current H5N1 vaccines and enhancing preparedness for future pandemics. Research should focus on elucidating the mechanisms of action underlying immune protection in the H5 influenza vaccine model, including the roles of adjuvants in recruiting both memory and naive B cell responses against conserved and strain-specific epitopes and the influence of adjuvants on antigen distribution and presentation. These efforts will be critical for guiding the rational design of next-generation vaccines with broad and durable efficacy against emerging H5 virus variants.

## Acknowledgments

DVR is supported by the U.S. Army’s Long-Term Health and Education Training Program. The following reagent was obtained through BEI Resources, NIAID, NIH: Peptide Array, Influenza Virus A/Vietnam/1203/2004 (H5N1) Hemagglutinin Protein, NR-18974.

## Author Contributions

Conceptualization: DVR, SD, MBF, RKE

Methodology: DVR, LB, SD, BMT, MBF, RKE

Investigation: DVR, LB, SD, BMT, SS, CD, VLS

Formal Analysis: DVR, BMT

Reagents (rHA): JH, SDV, HN, LC

Reagents (PR8/H5 virus): WS

Funding acquisition: DVR, MBF, RKE

Writing – original draft: DVR

Writing – review & editing: DVR, CD, MBF, RKE

All authors read and approved the manuscript.

## Data and materials availability

All data are available in the main text or the supplementary materials; further inquiries can be directed to the corresponding author.

## References

1 Treanor, J. J., Campbell, J. D., Zangwill, K. M., Rowe, T. & Wolff, M. Safety and immunogenicity of an inactivated subvirion influenza A (H5N1) vaccine. N Engl J Med 354, 1343–1351 (2006). 10.1056/NEJMoa055778

2 Treanor, J. J. et al. Safety and immunogenicity of a recombinant hemagglutinin vaccine for H5 influenza in humans. Vaccine 19, 1732–1737 (2001). 10.1016/s0264-410x(00)00395-9

3 Treanor, J. J. et al. Evaluation of safety and immunogenicity of recombinant influenza hemagglutinin (H5/Indonesia/05/2005) formulated with and without a stable oil-in-water emulsion containing glucopyranosyl-lipid A (SE+GLA) adjuvant. Vaccine 31, 5760–5765 (2013). 10.1016/j.vaccine.2013.08.064

4 Neuzil, K. M. et al. Safety and Immunogenicity of Influenza A/H5N8 Virus Vaccine in Healthy Adults: Durability and Cross-reactivity of Antibody Responses. Clin Infect Dis (2023). 10.1093/cid/ciac982

5 Winokur, P. L. et al. Safety and Immunogenicity of a monovalent inactivated influenza A/H5N8 virus vaccine given with and without AS03 or MF59 adjuvants in healthy adults. Clin Infect Dis (2023). 10.1093/cid/ciac983

6 Wiley, D. C. & Skehel, J. J. The structure and function of the hemagglutinin membrane glycoprotein of influenza virus. Annu Rev Biochem 56, 365–394 (1987). 10.1146/annurev.bi.56.070187.002053

7 Whittle, J. R. et al. Broadly neutralizing human antibody that recognizes the receptor-binding pocket of influenza virus hemagglutinin. Proc Natl Acad Sci U S A 108, 14216–14221 (2011). 10.1073/pnas.1111497108

8 Fischetti, L., Zhong, Z., Pinder, C. L., Tregoning, J. S. & Shattock, R. J. The synergistic effects of combining TLR ligand based adjuvants on the cytokine response are dependent upon p38/JNK signalling. Cytokine 99, 287–296 (2017). 10.1016/j.cyto.2017.08.009

9 Steinhagen, F., Kinjo, T., Bode, C. & Klinman, D. M. TLR-based immune adjuvants. Vaccine 29, 3341–3355 (2011). 10.1016/j.vaccine.2010.08.002

10 Goff, P. H. et al. Synthetic Toll-like receptor 4 (TLR4) and TLR7 ligands as influenza virus vaccine adjuvants induce rapid, sustained, and broadly protective responses. J Virol 89, 3221–3235 (2015). 10.1128/JVI.03337-14

11 Gregg, K. A. et al. A lipid A-based TLR4 mimetic effectively adjuvants a Yersinia pestis rF-V1 subunit vaccine in a murine challenge model. Vaccine 36, 4023–4031 (2018). 10.1016/j.vaccine.2018.05.101

12 Haupt, R. et al. Enhancing the protection of influenza virus vaccines with BECC TLR4 adjuvant in aged mice. Sci Rep 13, 715 (2023). 10.1038/s41598-023-27965-x

13 Haupt, R. E. et al. Novel TLR4 adjuvant elicits protection against homologous and heterologous Influenza A infection. Vaccine 39, 5205–5213 (2021). 10.1016/j.vaccine.2021.06.085

14 Das, S. et al. Enhancing protective efficacy and immunogenicity of hemagglutinin-based influenza vaccine utilizing adjuvants developed by BECC. bioRxiv, 2025.2006.2003.657703 (2025). 10.1101/2025.06.03.657703

15 Park, M. S., Steel, J., Garcia-Sastre, A., Swayne, D. & Palese, P. Engineered viral vaccine constructs with dual specificity: avian influenza and Newcastle disease. Proc Natl Acad Sci U S A 103, 8203–8208 (2006). 10.1073/pnas.0602566103

16 Hu, G. et al. Physicochemical characterization of biological and synthetic forms of two lipid A-based TLR4 agonists. Heliyon 9, e18119 (2023). 10.1016/j.heliyon.2023.e18119

17 Centers for Disease, C. & Prevention. FDA licensure of bivalent human papillomavirus vaccine (HPV2, Cervarix) for use in females and updated HPV vaccination recommendations from the Advisory Committee on Immunization Practices (ACIP). MMWR Morb Mortal Wkly Rep 59, 626–629 (2010).

18 Garcon, N., Segal, L., Tavares, F. & Van Mechelen, M. The safety evaluation of adjuvants during vaccine development: the AS04 experience. Vaccine 29, 4453–4459 (2011). 10.1016/j.vaccine.2011.04.046

19 Frasca, D., Diaz, A., Romero, M. & Blomberg, B. B. The generation of memory B cells is maintained, but the antibody response is not, in the elderly after repeated influenza immunizations. Vaccine 34, 2834–2840 (2016). 10.1016/j.vaccine.2016.04.023

20 Sasaki, S. et al. Limited efficacy of inactivated influenza vaccine in elderly individuals is associated with decreased production of vaccine-specific antibodies. J Clin Invest 121, 3109–3119 (2011). 10.1172/JCI57834

21 Frasca, D. et al. Effects of age on H1N1-specific serum IgG1 and IgG3 levels evaluated during the 2011-2012 influenza vaccine season. Immun Ageing 10, 14 (2013). 10.1186/1742-4933-10-14

22 Sun, X. et al. Broadly neutralizing antibodies to combat influenza virus infection. Antiviral Res 221, 105785 (2024). 10.1016/j.antiviral.2023.105785

23 Brandenburg, B. et al. Mechanisms of hemagglutinin targeted influenza virus neutralization. PLoS One 8, e80034 (2013). 10.1371/journal.pone.0080034

24 Guthmiller, J. J. et al. Broadly neutralizing antibodies target a haemagglutinin anchor epitope. Nature 602, 314–320 (2022). 10.1038/s41586-021-04356-8

25 Wu, N. C. & Wilson, I. A. Influenza Hemagglutinin Structures and Antibody Recognition. Cold Spring Harb Perspect Med 10 (2020). 10.1101/cshperspect.a038778

26 Zhang, Y. et al. A broad-spectrum vaccine candidate against H5 viruses bearing different sub-clade 2.3.4.4 HA genes. NPJ Vaccines 9, 152 (2024). 10.1038/s41541-024-00947-4

27 Luczo, J. M. & Spackman, E. Epitopes in the HA and NA of H5 and H7 avian influenza viruses that are important for antigenic drift. FEMS Microbiol Rev 48 (2024). 10.1093/femsre/fuae014

28 Bliss, C. M. et al. A single-shot adenoviral vaccine provides hemagglutinin stalk-mediated protection against heterosubtypic influenza challenge in mice. Mol Ther 30, 2024–2047 (2022). 10.1016/j.ymthe.2022.01.011

29 Wu, Y. et al. A potent broad-spectrum protective human monoclonal antibody crosslinking two haemagglutinin monomers of influenza A virus. Nat Commun 6, 7708 (2015). 10.1038/ncomms8708

30 Dreyfus, C. et al. Highly conserved protective epitopes on influenza B viruses. Science 337, 1343–1348 (2012). 10.1126/science.1222908

31 U.S. Food & Drug Administration, <https://www.fda.gov/vaccines-blood-biologics/influenza-vaccine-composition-2025-2026-us-influenza-season> (

32 Wang, W. et al. Serum Samples From Middle-aged Adults Vaccinated Annually with Seasonal Influenza Vaccines Cross-neutralize Some Potential Pandemic Influenza Viruses. J Infect Dis 213, 403–406 (2016). 10.1093/infdis/jiv407

33 Song, J. Y. et al. Randomized, double-blind, multi-center, phase III clinical trial to evaluate the immunogenicity and safety of MG1109 (egg-based pre-pandemic influenza A/H5N1 vaccine) in healthy adults. Hum Vaccin Immunother 13, 1190–1197 (2017). 10.1080/21645515.2016.1263410

34 Wong, S. S. et al. The immune correlates of protection for an avian influenza H5N1 vaccine in the ferret model using oil-in-water adjuvants. Sci Rep 7, 44727 (2017). 10.1038/srep44727

35 Lu, X. et al. A mouse model for the evaluation of pathogenesis and immunity to influenza A (H5N1) viruses isolated from humans. J Virol 73, 5903–5911 (1999). 10.1128/JVI.73.7.5903-5911.1999

36 Qiao, J. et al. Pulmonary fibrosis induced by H5N1 viral infection in mice. Respir Res 10, 107 (2009). 10.1186/1465-9921-10-107

37 Perrone, L. A., Plowden, J. K., Garcia-Sastre, A., Katz, J. M. & Tumpey, T. M. H5N1 and 1918 pandemic influenza virus infection results in early and excessive infiltration of macrophages and neutrophils in the lungs of mice. PLoS Pathog 4, e1000115 (2008). 10.1371/journal.ppat.1000115

38 Lazniewski, M., Dawson, W. K., Szczepinska, T. & Plewczynski, D. The structural variability of the influenza A hemagglutinin receptor-binding site. Brief Funct Genomics 17, 415–427 (2018). 10.1093/bfgp/elx042

39 Russell, R. J. et al. H1 and H7 influenza haemagglutinin structures extend a structural classification of haemagglutinin subtypes. Virology 325, 287–296 (2004). 10.1016/j.virol.2004.04.040

40 Wang, W. et al. Human antibody 3E1 targets the HA stem region of H1N1 and H5N6 influenza A viruses. Nat Commun 7, 13577 (2016). 10.1038/ncomms13577

41 DiLillo, D. J., Tan, G. S., Palese, P. & Ravetch, J. V. Broadly neutralizing hemagglutinin stalk-specific antibodies require FcgammaR interactions for protection against influenza virus in vivo. Nat Med 20, 143–151 (2014). 10.1038/nm.3443

42 Mokalla, V. R., Gundarapu, S., Kaushik, R. S., Rajput, M. & Tummala, H. Influenza Vaccines: Current Status, Adjuvant Strategies, and Efficacy. Vaccines (Basel) 13 (2025). 10.3390/vaccines13090962

43 Ellebedy, A. H. et al. Adjuvanted H5N1 influenza vaccine enhances both cross-reactive memory B cell and strain-specific naive B cell responses in humans. Proc Natl Acad Sci U S A 117, 17957–17964 (2020). 10.1073/pnas.1906613117

44 Griffin, M. R. et al. Effectiveness of non-adjuvanted pandemic influenza A vaccines for preventing pandemic influenza acute respiratory illness visits in 4 U.S. communities. PLoS One 6, e23085 (2011). 10.1371/journal.pone.0023085

45 Ray, G. T. et al. Intraseason Waning of Influenza Vaccine Effectiveness. Clin Infect Dis 68, 1623–1630 (2019). 10.1093/cid/ciy770

46 Young, B., Sadarangani, S., Jiang, L., Wilder-Smith, A. & Chen, M. I. Duration of Influenza Vaccine Effectiveness: A Systematic Review, Meta-analysis, and Meta-regression of Test-Negative Design Case-Control Studies. J Infect Dis 217, 731–741 (2018). 10.1093/infdis/jix632

